# A Polygenic Quincunx

**DOI:** 10.1101/2023.07.18.549486

**Authors:** Ralph H. Stern

**Affiliations:** Division of Cardiovascular Medicine, Department of Internal Medicine

## Abstract

Sequential addition of SNPs (single nucleotide polymorphism) to a risk model repeatedly splits risk groups in two, generating a branching tree. As SNPs are independent, the path of an individual through the tree is a form of random walk. This resembles the path of a ball through a Galton quincunx, but the rules for a polygenic quincunx are more complex. The result is that individuals zigzag to higher and lower risks with each additional SNP. Patient flows through a polygenic quincunx for a few SNPs that aggregate these individual random walks is presented. Longer random walks calculated from Monte-Carlo simulations illustrate how different the individual random walks can be. As random walks do not have unique termini, polygenic risk values don’t converge on unique individual risk estimates. However, the more steps in a random walk, the greater the dispersion of the termini around the starting point. So more SNPs can result in more disperse population polygenic risk distributions, and potentially greater clinical benefit when allocating preventive measures based on risk level.

---

Polygenic risk distributions for a population are well described in the literature, but the fates of individuals are not. Sequential addition of SNPs (single nucleotide polymorphisms) to a risk model repeatedly splits risk groups in two, generating a branching tree. As SNPs are independent, the path of an individual through the tree is a form of random walk. This resembles the path of a ball through a Galton quincunx, an array of offset pegs. When a ball is dropped above the array in the middle, it bounces off a peg in the first layer: half of the time to the left and half of the time to the right. As it falls further, this is repeated at each subsequent layer of pegs with the ball zigzagging right and left at random. The distribution of balls after the last layer of pegs is approximated by the binomial distribution. In this paper the more complex rules for a polygenic quincunx are utilized to simulate the random walks of individuals on addition of SNPS to risk models. That individuals follow a random walk on the addition of SNPs has implications for the interpretation of polygenic risk for individuals.

## METHODS

There are two methods for simulating the results of sequential addition of SNPs: calculation from the binomial distribution of the expected results or Monte Carlo simulation of random walks. Each gene contributes two independent SNPs.

For the first method, risk alleles are assumed to have the same prevalence and relative risk or odds ratio and to interact multiplicatively. Assuming Hardy-Weinberg equilibrium and independent loci, the distribution of risk alleles is given by the binomial distribution, (q+p)^n^ (n=number of SNPs, p=prevalence of risk allele, q=(1-p)=(1-prevalence of risk allele)).^1^

The use of an odds ratio model avoids calculating absolute risks above one.^2^ For an odds ratio (OR) model, the distribution of odds ratios is given by: (q+p OR)^n^ and the distribution of odds is given by: O_0_ (q+p OR)^n^, where O_0_ is the odds of the group with zero risk alleles specific for that n. Solving for O_0_ requires converting odds to absolute risks to generate the weighted sum of absolute risks, which equals the overall risk (R), and then solving numerically, which does not require Monte Carlo simulation:

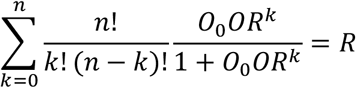

This method was used to calculate polygenic risk distributions for different values of n.

For a relative risk model, the distribution of relative risks (RR) is given by: (q+p RR)^n^ and the distribution of absolute risks by: R_0_ (q+p RR)^n^, where R_0_ is the absolute risk of the group with zero risk alleles specific for that n. Since the latter expression is the weighted sum of absolute risks, it is equal to the overall risk (R), which can be either the population risk or the risk of a specific risk stratum:

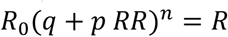

This can be solved for R_0_, which does not require Monte Carlo simulation. This method was used for creating a polygenic quincunx. The risks in each layer of the quincunx are R_0_, R_0_ RR, R_0_ RR^2^, …, R_0_ RR^n^.

For longer random walks, Monte Carlo simulation was employed to calculate changes in risk as

SNPs are sequentially added. As above, starting at a risk of R, which can represent the population risk or the risk of a specific risk stratum, addition of a SNP splits the population or risk stratum in two: a higher risk fraction p with the risk allele and a lower risk fraction q=1-p without the risk allele. The weighted sum of risks of the two risk subgroups must equal R, allowing solution for the risks in the two subgroups. For an odds ratio model, the risk of the lower risk subgroup is:

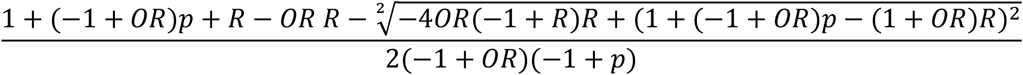

and the risk of the higher risk subgroup is:

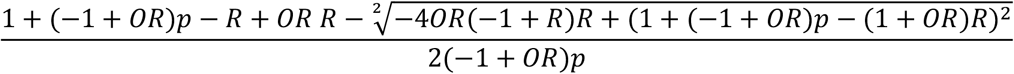

As more SNPs are added, the results of the subsequent splits can in turn be calculated. Five random walks generated by adding SNPs (p 0.2, OR 1.1) starting at R=0.1 were simulated.

The Monte-Carlo simulations do not produce exactly the same risks as those of the binomial distribution. In the Monte-Carlo simulation, risks are not constrained to be those given by the binomial distribution (where there is a single O_0_ or R_0_ per layer and the risk sub groups are obtained by multiplying by powers of OR or RR, respectively, whereas in a random walk there is an O_0_ or R_0_ calculated for each split) Instead, a narrow distribution of risks around the binomial risks are obtained, but the weighted sum of these risks remains equal to R.

## RESULTS

An example of probability mass functions and cumulative distribution functions for polygenic risk distributions is shown in Figure 1 for R=0.1, p=0.2, and OR=1.1 and n=100, 1,000, and 10,000 SNPs. As the number of SNPs is increased, the risk distributions become more disperse and more skewed.

**Figure 1.**
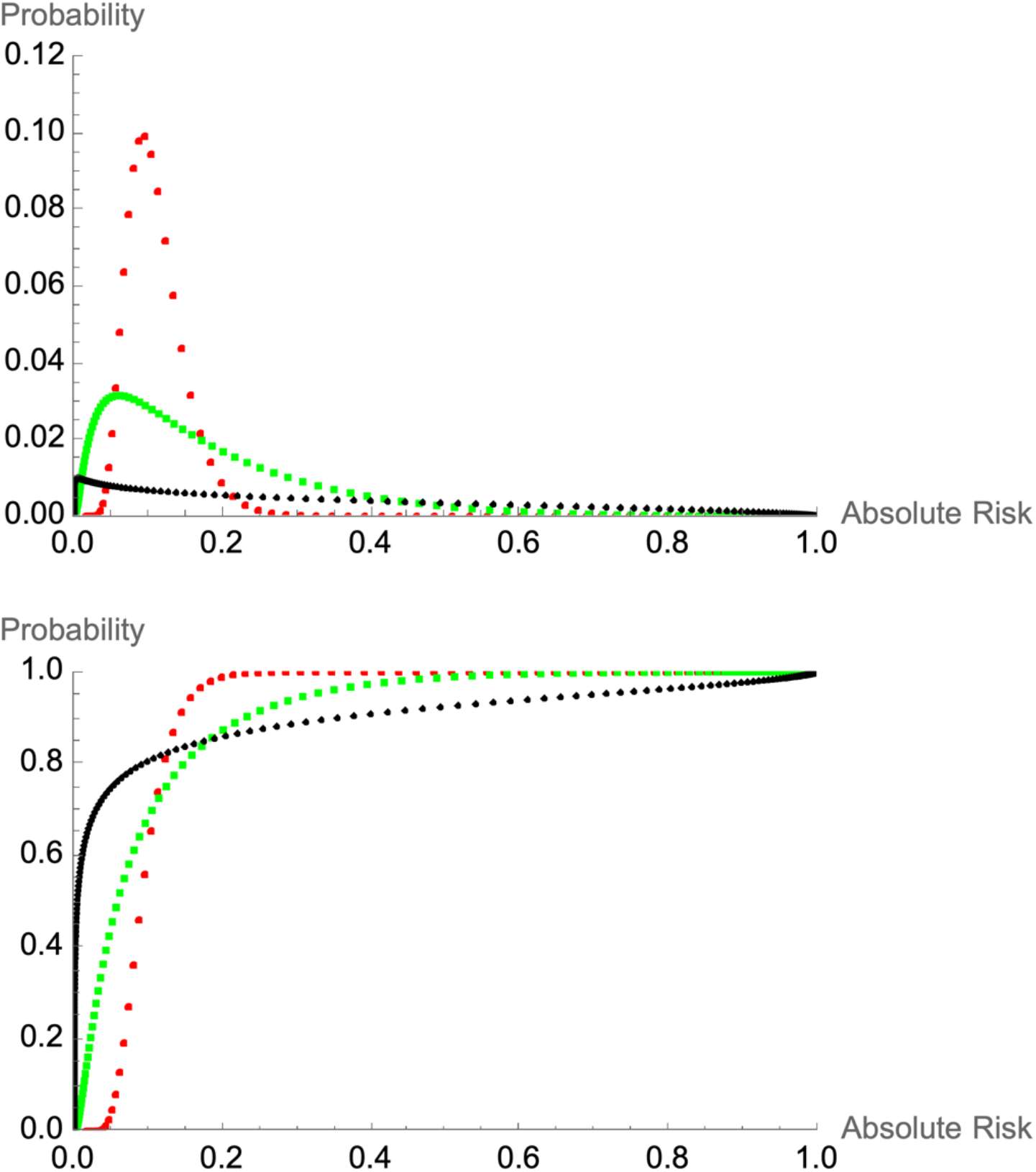
Probability mass functions (above) and cumulative distribution functions (below) for polygenic risk distributions for R=0.1, p=0.2, and OR=1.1 and n=100 (red), 1,000 (green), and 10,000 (black) SNPs

An interesting incidental observation is that two risk distributions may place the same fraction of the population above a selected risk, but the risks of that fraction may differ considerably. Both the 1,000 and 10,000 SNP models place about 0.15 of the population above an absolute risk of 0.2, but this high-risk group is much riskier when identified by the 10,000 SNP model. This would be missed by metrics that only assess differences in the number of patients in a risk category.

For the quincunx, the splitting of risk groups on addition of a SNP is presented in Figure 2. Addition of a SNP splits the population or risk stratum in two: a higher risk fraction p with the risk allele and a lower risk fraction q=1-p without the risk allele. As a specific example, patient flows thru a polygenic quincunx with 6 layers (corresponding to sequential addition of 6 SNPs) starting at R=0.1 with RR=1.1 and p=0.2 for 100,000 subjects are shown in Figure 2. Since relative risks were used, this same distribution would be observed starting at any R (unless risks above 1 were to result), so relative risks rather than absolute risks are presented on the risk axis.

**Figure 2.**
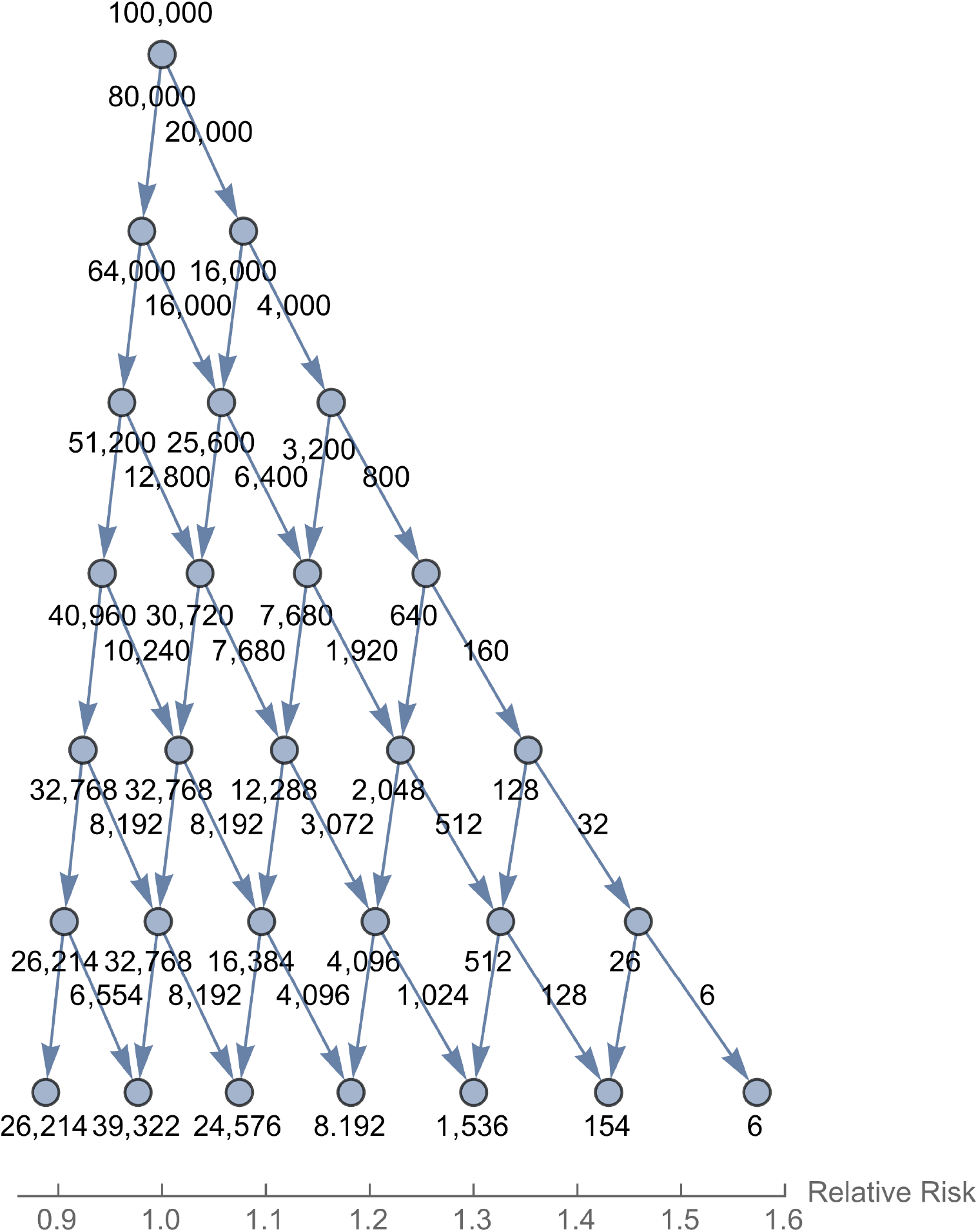
Patient flow of 100,000 individuals through a polygenic quincunx for R=0.1, p=0.2, and OR=1.1

Examples of random walks with 500 steps (corresponding to addition of 500 SNPs or 250 genes) for five individuals are shown in Figure 3. These were derived from Monte Carlo simulations, starting at R=0.1 with OR=1.1 and p=0.2

**Figure 3.**
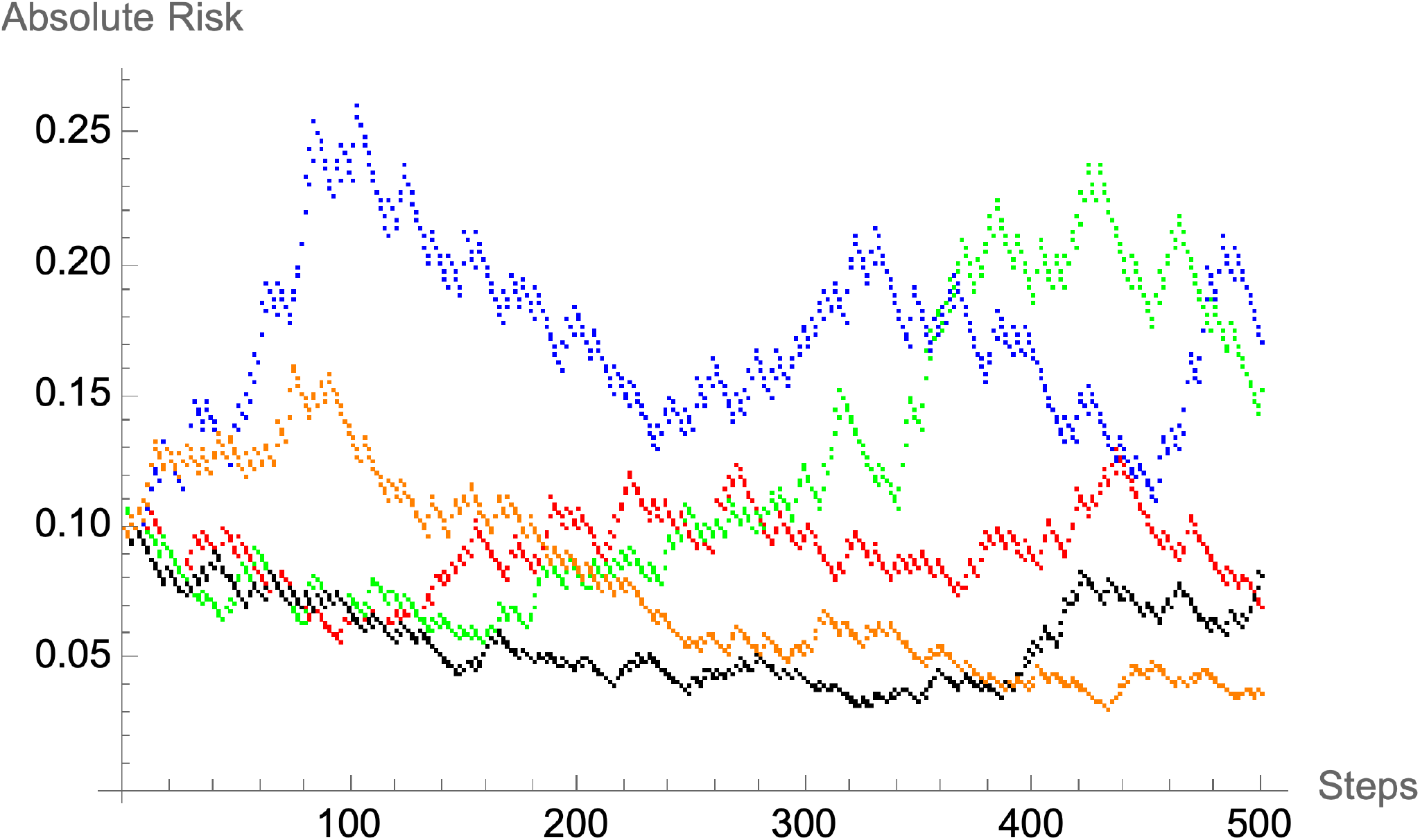
Five random walks of 500 steps for p=0.2 and OR=1.1 starting at R=0.1

## DISCUSSION

For additions to a risk model to improve clinical outcomes for a population, they need to improve risk stratification, i.e., increase the dispersion of risk subgroups around the population risk, as this may allow improvements in selective allocation of preventive measures (treatments or surveillance) based on risk.

Subgroups with lower risk might not merit the same preventive measures as those at average risk, while those with higher risk might merit more intensive or extensive preventive measures than those at average risk. For polygenic risk, the greater the number of SNPs, the more disperse the polygenic risk distribution and the greater the potential clinical benefit. The more disperse the polygenic risk distribution, the greater the separation of patient and nonpatient risks, which is quantified by measures of discrimination, the ROC curve AUC and c-index. For the polygenic risk distributions in Figure 1, the ROC curve AUCs are 0.606, 0.788, and 0.964 for n=100, 1,000, and 10,000, respectively. Improvements in population risk stratification of this magnitude would be expected to lead to superior ability to allocate preventive measures.

Assuming the risks are accurate (calibrated), this increasing dispersion cannot be achieved by simply expanding the risk distribution like an accordion; the rank order of individuals must also change. For example, a group of individuals at 15% risk cannot simply be shifted to risks between 15 and 20% if the 15% risk is accurate. This means there must be continual shuffling of the rank order of individuals as SNPs are added.

This shuffling of individuals can be seen in the quincunx and random walks in Figures 2 and 3. As SNP’s are added, individual subjects zigzag back and forth at random to lower and higher risks without homing in on a unique individual risk. This can be observed even when addition of SNPs does not noticeably improve dispersion and discrimination measures.^3^ If the continuous range of absolute risks is categorized into treatment groups, these movements can result in individuals crossing a category boundary, referred to as reclassification. And sequential addition can lead to individuals passing back and forth across these boundaries and receiving alternating treatment recommendations.^4^

Kaplan and Turkheimer have pointed out the similarities between genome wide association studies and Galton’s quincunx in their discussion of causation in behavior genetics.^5^ Galton’s quincunx used equally spaced pegs so the distribution of balls after the last layer of pegs approaches the binomial distribution and is approximated by the continuous normal distribution. An alternative design has an array of triangular shapes instead of pegs. These are constructed so that the steps to right or left are multiplicative instead of additive. The distribution of balls after the last layer is approximated by the lognormal distribution.^6^ A virtual polygenic quincunx differs from these physical quincunxes in that steps are not simply additive (stepping up from R to R+constant or stepping down from R to R-constant) or multiplicative (stepping up from R to R*constant or stepping down to R/constant) and the splitting is not symmetric.

If SNPs were added in order of decreasing OR or RR until the maximum number of SNPs available was reached, it might give the appearance of individuals converging on a single risk value, but this would not represent homing in on a unique individual risk.

As random walks do not have unique termini, polygenic risk scores should not be interpreted as estimates of an individual’s risk (even were SNPs the only risk factors known), nor should addition of SNPs be interpreted as leading to more precise estimates of an individual’s risk. Separate from these considerations, there is good reason to doubt that individual risk exists.^7^ But adding steps to a random walk does lead to a greater dispersion of the termini around the starting point. And, as discussed above, the more disperse the polygenic risk distribution, the greater the potential clinical benefit.

## Notes

### Competing Interest Statement

The authors have declared no competing interest.

